# Multi-platform reassessment of human mitochondrial DNA methylation reveals signals consistent with technical artifacts

**DOI:** 10.64898/2026.06.10.730935

**Authors:** Salman Basrai, Alexander T Bahcheli, Djarren Tan, Philip C Zuzarte, Andrea Bevan, Theodore Chan, Kristie Ng, Bernard Lam, Andrea Arruda, Sunit Das, Mark D Minden, Jared T Simpson, Jüri Reimand, Sagi Abelson

## Abstract

The existence and functional relevance of mitochondrial DNA methylation remain controversial. Here, we systematically profiled cytosine methylation and hydroxymethylation across human brain and blood tissues spanning healthy and malignant states using orthogonal sequencing approaches that avoid chemical conversion during library preparation. While nuclear DNA exhibited canonical methylation patterns, mitochondrial DNA consistently showed negligible signal, indistinguishable from background technical noise. By mapping cytosine-guanine sites between mitochondrial DNA and nuclear-embedded mitochondrial sequences, we demonstrate the potential of these nuclear counterparts to confound not only cytosine methylation but also hydroxymethylation measurements, corroborating and extending prior findings implicating nuclear contamination as a potential source of apparent mitochondrial epigenetic signals. Additional technical factors that inflate apparent mtDNA methylation signals were identified, including sequence context biases, flow cell chemistries, and coverage-dependent discrepancies between the heavy and light strands. Collectively, these results provide convergent evidence against the presence of biologically meaningful cytosine methylation or hydroxymethylation in mitochondrial DNA. These findings caution against interpreting apparent mtDNA methylation signals in human adult tissues as meaningful without rigorous orthogonal validation and comprehensive consideration of technical and analytical confounding factors.

## Background

Human mitochondrial DNA (mtDNA) is a compact, circular genome essential for cellular energy production.^1^ Historically, cytosine methylation, a key epigenetic modification modulating nuclear gene expression and genome stability, was considered absent from mtDNA. This view arose from fundamental differences between the nuclear and mitochondrial genomes, including their distinct structures, separate cellular compartments, and the lack of canonical epigenetic regulator genes encoded in the mitochondrial genome. These distinctions extended to sequence context: nuclear DNA methylation predominantly occurs at CpG dinucleotides,^2^ whereas the majority of cytosines in mtDNA (∼88%) are found in non-CpG (CpH) contexts.^3^ Given the relatively low frequency of CpH methylation in most differentiated tissues and the predominantly nuclear localization of DNA methylation machinery, these observations further argued against widespread mtDNA methylation in adult human somatic cells under physiological conditions.^4^ Consistent with this view, early investigations using semi-quantitative approaches failed to detect 5-methylcytosine (5mC) in mtDNA, reinforcing the notion that mtDNA methylation is minimal or absent.^5^

This long-standing paradigm has been challenged by accumulating reports suggesting that human mtDNA may carry cytosine modifications. Advances in genome-wide and enrichment-based detection technologies, including methylated DNA immunoprecipitation (MeDIP) and bisulfite sequencing, have reported low-level methylation-like signals within the human mitochondrial genome across a range of tissues and cell types. These studies have suggested the presence 5mC and, in some cases, 5-hydroxymethylcytosine (5hmC) within mtDNA, with proposed associations to mitochondrial function and disease states.^6^ Corroborating these findings, multiple studies have proposed the mitochondrial localization and import of DNA methyltransferases (DNMTs) and Ten-eleven translocation (TET) isoforms, raising the possibility of enzymatic activity contributing to catalyzing 5mC and 5hmC modifications in mtDNA.^7–9^ Literature reviews and accumulating reports continue to describe dynamic and locus-specific 5mC and 5hmC patterns in mtDNA, with proposed biological relevance and potential applications as biomarkers and exploratory therapeutic targets.^10,11^

Nonetheless, a major point of contention remains the lack of reproducible, independent and orthogonal evidence linking proposed mtDNA methylation to definitive functional consequences under physiological or pathological conditions. Concerns have also been raised regarding potential technical artifacts, including incomplete bisulfite conversion, nuclear DNA carryover, and inadequate technical or biological controls. Consequently, the existence, extent, and biological significance of mtDNA methylation remain highly contentious. This debate is perhaps best exemplified by two studies that analyzed the same human DNA methylation datasets yet reached opposing conclusions, underscoring the persistent uncertainty surrounding the field.^12,13^

Resolving whether mtDNA methylation occurs at appreciable, functionally relevant levels is crucial. If substantiated, mtDNA methylation could represent a distinct layer of epigenetic regulation that potentially modulates mitochondrial gene expression and function, influences cellular metabolism, and contributes to the pathogenesis of various diseases.^6^ Otherwise, much of the research in this area remains vulnerable to scrutiny, underscoring the need for particularly stringent evidentiary standards for the validation and interpretation of future mtDNA methylation claims in the scientific literature.

To address the ongoing controversy in the field and technical uncertainties surrounding mtDNA methylation detection, we employed a multi-platform strategy combining distinct library preparation approaches and sequencing technologies. Three platforms were used: enzymatic multi-omic library preparation coupled to Illumina sequencing using the Biomodal platform^14^, sequencing of native DNA using Oxford Nanopore technology, which provides direct signal-level inference of DNA molecules, and enzymatic methylation sequencing (EM-seq) library preparation coupled with Ultima Genomics proprietary sequencing.^15^ Together, these approaches encompass short- and long-read sequencing, native DNA sequencing and enzymatic conversion-based methylation profiling, and detection of both 5mC and 5hmC across orthogonal platforms.

To investigate tissue-specific variation and determine whether elevated nuclear DNA methylation also reflect in robust mtDNA methylation signals, we analyzed human brain tissue, and mobilized blood stem cells characterized by high methylation levels^2,16^. Lastly, to contextualize mtDNA methylation in pathological conditions, we analyzed samples from patients with glioblastoma (GBM) and acute myeloid leukemia (AML), a hematologic cancer characterized by frequent mutations in epigenetic regulator genes and well-established epigenetic dysregulation^17^.

## Results

### Widespread nuclear epigenetic divergence between brain and blood is not reflected in mitochondrial DNA

Using Biomodal sequencing,^14^ we quantified methylation levels across 158,823 non-overlapping 16,569 bp windows tiling the nuclear genome, selected to match the length of mtDNA. In comparing brain and blood 62,389 windows (39.28%) exhibited at least a twofold increase in 5mC levels in the brain, while 7,561 windows (4.76%) showed at least a twofold increase in 5hmC levels (two-sided Wilcoxon rank-sum test; Bonferroni corrected P < 0.05). In addition, 400 windows (0.25%) exhibited at least a twofold enrichment in both 5mC and 5hmC marks. Conversely, only eight windows displayed statistically significant methylation enrichment in blood compared to brain (Table S1).

Genome-wide visualization revealed that differential 5mC was broadly distributed across the nuclear genome, typically showing moderate fold changes between tissues. In contrast, differential 5hmC was largely confined to open chromatin regions and frequently associated with higher fold-change magnitudes (Figure S1 and Figure S2). These patterns are consistent with the known enrichment of nuclear DNA methylation in brain tissue and the functional roles of 5mC and 5hmC in chromatin architecture and gene regulation^2,18^, thereby providing internal support for the analytical framework employed.

In stark contrast to the distinct nuclear methylation landscapes revealed by this window-based tiling analysis, the mitochondrial genome showed no statistical difference in global DNA methylation levels between brain and blood samples. This finding was consistent across analytical resolutions, as differential methylation analyses performed without restriction to fixed windows likewise failed to identify any differentially methylated regions. At base resolution, mtDNA profiles were highly concordant across samples and characterized by uniformly low levels of both 5mC and 5hmC, consistent with residual signal at the level of technical background of the Biomodal approach^14^ (Figure 1).

**Figure 1.**
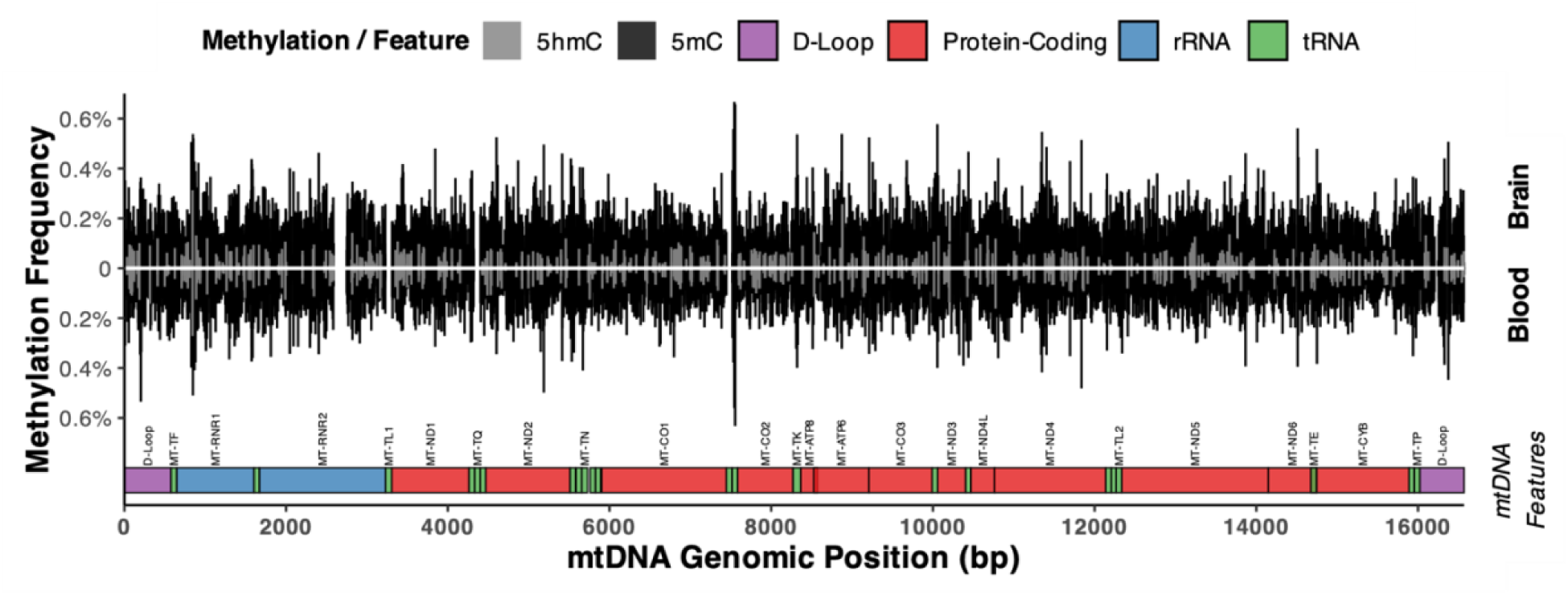
Distribution of low-level 5-methylcytosine (5mC) and 5-hydroxymethylcytosine (5hmC) signal across human mitochondrial DNA (mtDNA). The upper panel shows the brain mtDNA methylation profile (positive y-axis), and the lower panel shows the blood profile (negative y-axis). Methylation frequency (%) is shown for detected sites. The bottom track annotates mtDNA features, including protein-coding genes (red), rRNA genes (blue), tRNA genes (green), and the D-loop (purple), with feature names indicated vertically. Right-hand labels indicate tissue assignment and feature track position. Gaps correspond to soft-masked regions of the mtDNA reference sequence, representing low-complexity or repetitive elements.

### Trinucleotide sequence-context reveals artifactual origin of low-level mtDNA CpH methylation signals

mtDNA methylation have been reported to be enriched in non-CpG (CpH) contexts^12,19,20^. However, base substitution errors in deep sequencing data are strongly context dependent.^21,22^ Drawing upon our prior characterization of sequencing error profiles^22^, we evaluated the apparent signals detected in the brain and blood mitochondrial genomes. Specifically, we examined methylation patterns within trinucleotide contexts and identified a distinct signature at CpH sites (Figure 2). To assess whether this pattern could arise from technical artefact, we performed a computational simulation in which nuclear genome-derived reads, which are expected to lack appreciable CpH methylation,^4^ were randomly subsampled to match the coverage and sequence-context distribution observed in mtDNA. These results demonstrate that the resulting pattern closely recapitulates the apparent low-level mtDNA CpH signal, suggesting that the inferred methylation reflects background noise inherent to the sequencing method and analytical framework rather than genuine biological modification.

**Figure 2.**
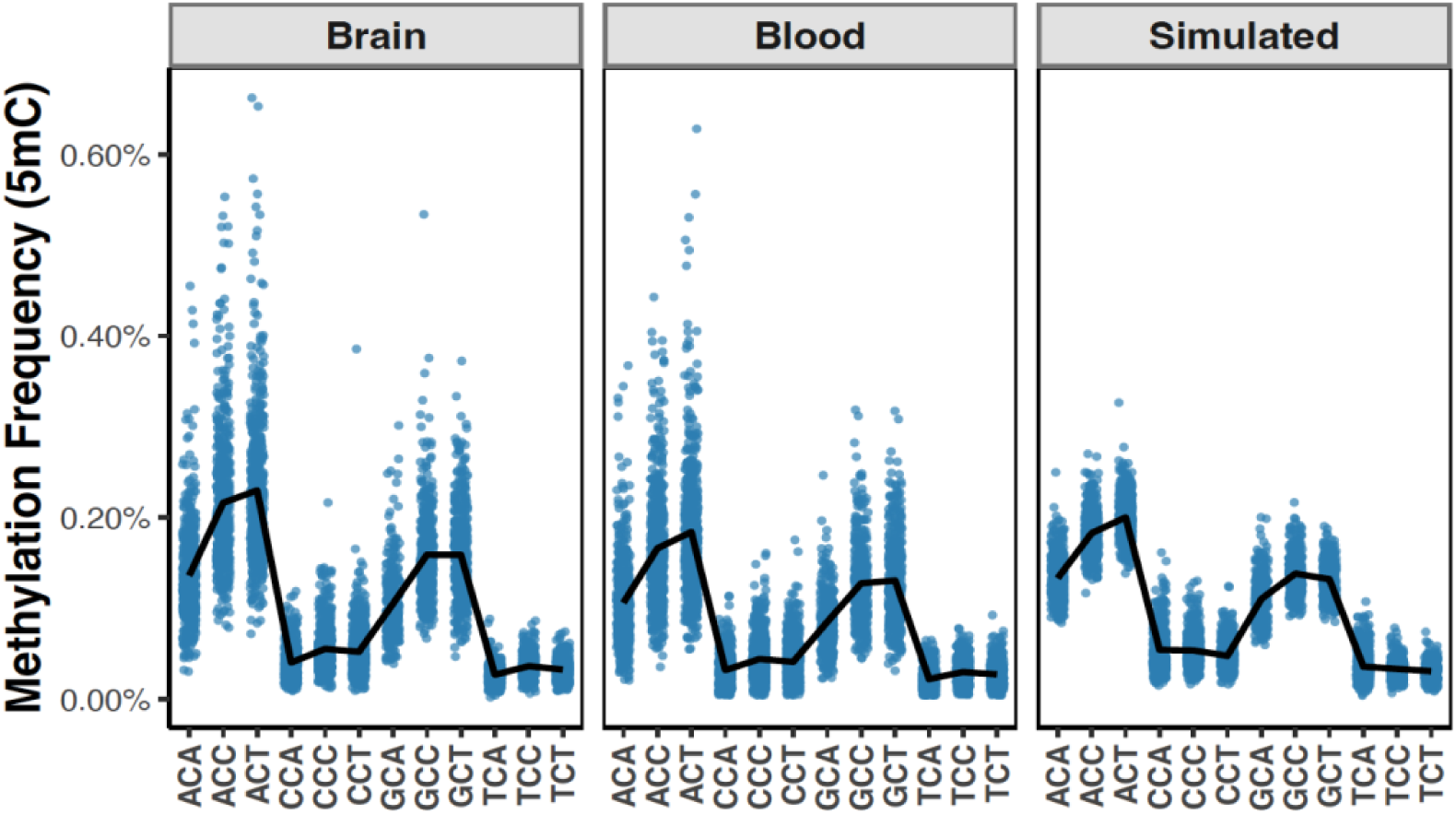
CpH methylation patterns and simulation-based assessment. Comparison of 5mC frequency in CpH trinucleotide contexts across brain, blood, and simulated mtDNA data derived from nuclear reads matched for sequence context and coverage. Each blue point represents methylation frequency at an individual CpH site (n = 4,641). The black line connects the mean 5mC frequency for each CpH context within each sample. X-axis labels indicate CpH trinucleotide contexts. Panels share a common y-axis.

### Long-read sequencing implicates nuclear-integrated mitochondrial sequences as a potential source of mtDNA 5mC and 5hmC signals, while revealing no appreciable mitochondrial methylation in cancer with mutations in epigenetic regulators

Recognizing epigenetic dysregulation as a cancer hallmark^23^ that may extend to mitochondria, we investigated mtDNA methylation in brain and blood malignancies using Nanopore sequencing. This platform enables direct detection of DNA modifications without chemical or enzymatic cytosine conversion, thereby avoiding technical biases associated with conversion-based methylation profiling approaches.^24^

Our analysis focused on quantifying 5mC and 5hmC within the CpG dinucleotide context, as the lower signal-to-noise ratio currently limits reliable detection of CpH methylation by Nanopore sequencing.^25^ Long-read sequencing was performed on samples from patients with two distinct malignancies, glioblastoma (GBM) and acute myeloid leukemia (AML). GBM was selected based on prior bisulfite sequencing report suggesting elevated mtDNA methylation.^26^ AML cases were selected to represent prevalent epigenetic subtypes, with each patient carrying mutations in a major epigenetic regulator: one sample harbors two nonsense mutations in *TET2*, another sample was detected with two missense mutations in *DNMT3A*, and the third has an *IDH2* hotspot mutation known to drive widespread DNA methylation changes (Table S2, Figure S3).^27–29^ As a negative control for methylation detection, we also analyzed whole genome amplified DNA, a process which removes existing epigenetic marks. To establish a robust positive internal control for 5mC and 5hmC detection, we leveraged nuclear-integrated mitochondrial sequences (NUMTs) which are mitochondrial-derived DNA fragments integrated into the nuclear genome during evolution and therefore subject to canonical nuclear DNA methylation.^30^ The high sequence homology between NUMTs and mtDNA ensures comparable electrical signal profiles, minimizing sequence-context bias when comparing CpG modification calls between reads originating from the nuclear genome versus the mitochondrial genome.

Across all patient samples analyzed, irrespective of tissue of origin or underlying genomic alterations, mtDNA methylation exhibited negligible levels, consistent with background signal (Table S3). The 5mC rate - calculated as the proportion of modified cytosines relative to the total number of sequenced cytosines - was zero in the negative control sample. In contrast, the 5hmC rate in the same sample was 0.0117%, corresponding to approximately one modified cytosine per 10,000 sites interrogated, and was the second highest among all samples.

Analysis of 796 confidently mapped mitochondrial CpG sites within NUMTs (91.49% of all 870 mtDNA CpGs) revealed significantly higher methylation levels in the nuclear genome compared to the corresponding mitochondrial loci for both 5mC and 5hmC across patient samples (two-sided Wilcoxon signed-rank test; P = 0.036), consistent with the expected higher methylation state of nuclear DNA (Figure 3, Table S4, Figure S4).

**Figure 3.**
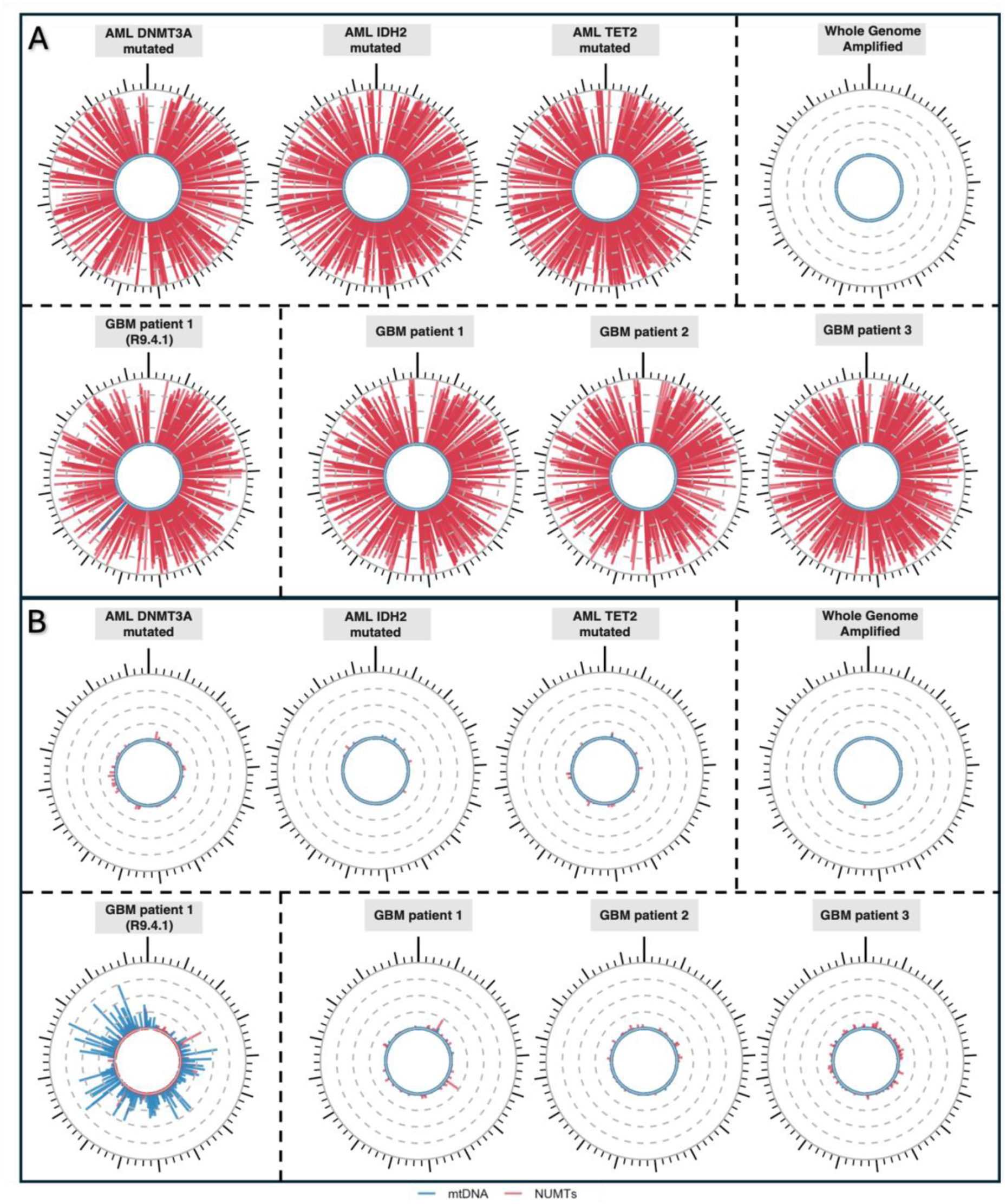
Position-matched comparison of methylation frequencies between mtDNA and nuclear-embedded mitochondrial DNA sequences (NUMTs). Circular plots visualize 5mC **(A)** and 5hmC **(B)** frequencies across the entire mitochondrial genome (16,569 bp). Each bar corresponds to a single CpG site and is colored by origin: mitochondrial DNA observations in blue and NUMTs in red. mtDNA values are plotted above position-matched NUMT values. Bar heights correspond to methylation frequency, scaled from 0 (unmethylated) to 1 (fully methylated; full radius). Gray guide circles indicate 0.25 methylation increments. Kilobase and 200 bp markers are displayed radially, with the first base positioned at the top. Only positions with at least 10x read coverage are shown.

Overall, these findings argue against appreciable mtDNA methylation in human cancer tissues and identify germline NUMTs as a potential source of falsely elevated mtDNA 5mC and 5hmC signals across most mitochondrial CpG sites.

### Nanopore flow cell version impacts inferred mtDNA methylation levels

To assess the impact of technological advancements on mtDNA methylation levels, we compared Nanopore sequencing data from the same GBM (patient 1) DNA sample, generated using two different flow cell versions. The earlier version (R9.4.1) yielded substantially higher mtDNA CpG methylation rates - 0.1757% for 5mC and 3.68% for 5hmC - compared to the more recent version (Table S3). These findings are markedly higher, by over one to nearly three orders of magnitude, than the negligible levels detected using the higher-accuracy platform employed in our main study (R10.4.1, released in Q1 2023; Figure 3). Given the known advancements in pore chemistry and base modification calling on the nanopore platform, this discrepancy underscores how technical variability in earlier sequencing workflows may have contributed to previous reports of detectable mtDNA methylation^6^ - findings that may not hold with state-of-the-art methods.

### Apparent mtDNA methylation signals are explained by extreme light-heavy strand coverage asymmetry and associated low local sequence complexity

We used Ultima Genomics sequencing as an orthogonal approach to assess mtDNA cytosine modification levels. Given prior reports of elevated CpG and non-CpG nuclear DNA methylation in hematopoietic stem and progenitor cells relative to differentiated populations,^16^ a mobilized peripheral blood stem cell sample was selected to assess whether analogous cytosine modification signals could also be detected in mtDNA within a stem-cell-enriched cellular context. EM-seq was used to quantified 5mC frequencies across the CpG, CHG, and CHH sequence contexts.

Across all sequence contexts, methylation frequencies exhibited a pronounced decay with increasing coverage (Exponential decay fit: CpG: R² = 0.764, CHG: R² = 0.757, CHH: R² = 0.766). This strong inverse relationship suggests that apparent methylation signals are substantially influenced by technical biases. To distinguish potential biological methylation from assay background, we analyzed unmethylated Lambda NEB phage DNA as a negative control. The Lambda control demonstrated low global error rates that were broadly consistent across contexts (range: 0.29%–0.31%). Across all contexts, 99% of error-associated methylation frequencies were detected below 1.78%, although occasional values reached as high as 15% (Figure 4A). Notably, the asymptotic methylation frequencies estimated for mtDNA at high coverage converged to values comparable to the global error rates observed in the Lambda control, indicating a shared lower bound defined by the intrinsic error characteristics of the assay. Methylation signal and coverage differed between the Heavy and Light strands, with significantly lower coverage and higher apparent methylation levels in the light strand (Figure 4B).

**Figure 4.**
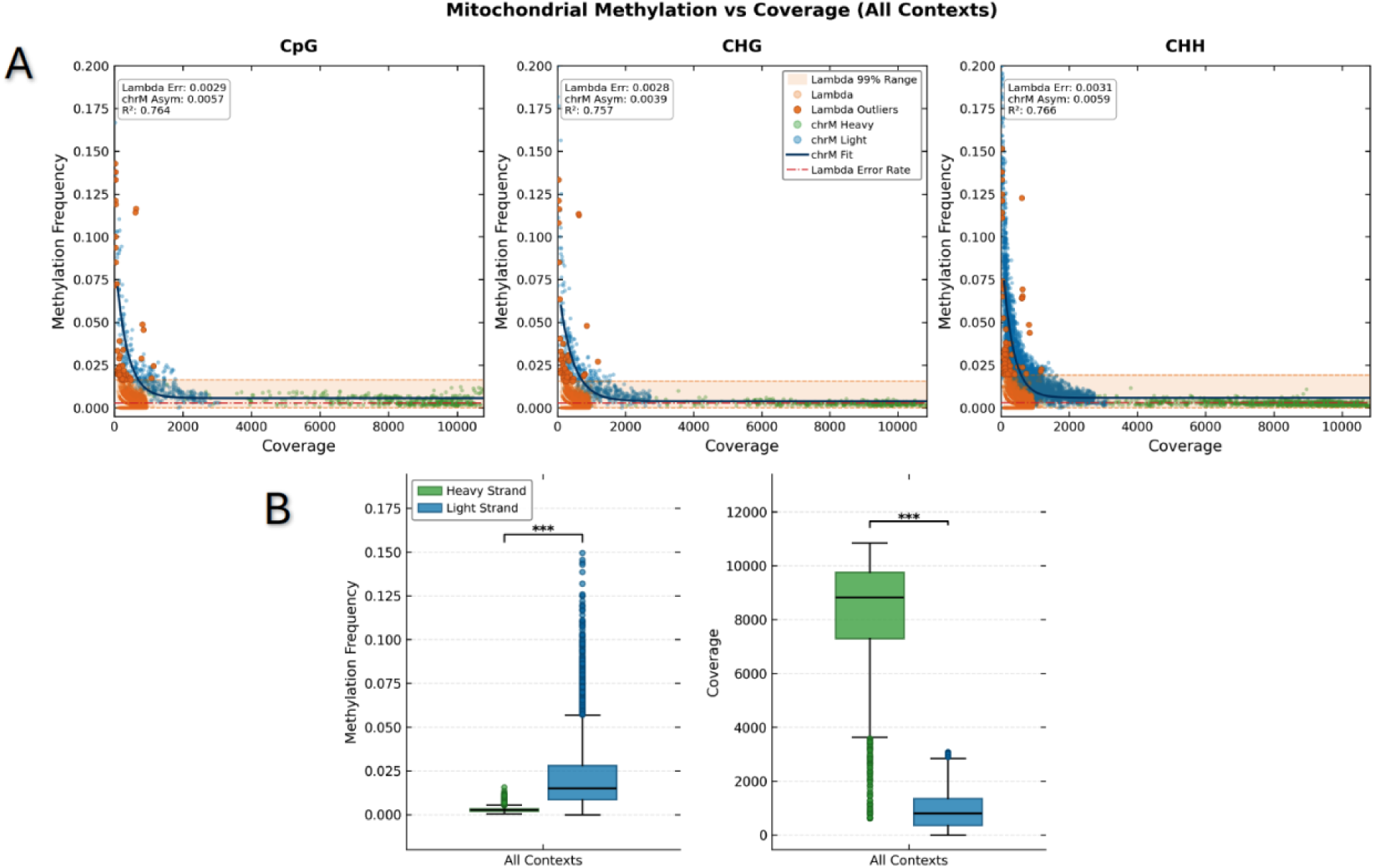
Coverage-dependent methylation decay and strand-specific biases in mitochondrial DNA. **(A)** Scatter plots depict the relationship between sequencing coverage and apparent methylation frequency for mtDNA and a lambda DNA negative control across CpG, CHG, and CHH sequence contexts. Each point represents a single cytosine. Lambda control data (light orange) define the technical error floor, with the 99% confidence range shaded and outliers above that interval highlighted in dark orange. Mitochondrial data are stratified by DNA strand: Heavy (green) and Light (blue). A black curve represents an exponential decay model fitted to mtDNA data, capturing the inverse relationship between coverage depth and apparent methylation. The red dash-dot line indicates the global lambda error rate. **(B)** Boxplots compare methylation frequencies (left) and sequencing coverage distributions (right) between the Heavy and Light mitochondrial strands, aggregated across all sequence contexts. Methylation values are restricted to sites with coverage ≥ 50x to minimize low-coverage stochastic noise. Statistical significance was assessed using the two-sided Mann-Whitney U test. Asterisks denote significance levels: ***p < 0.001.

A focused analysis of the light strand revealed that methylation signals, irrespective of sequence context, scaled inversely with coverage (Table S5, Figure 5). Sequencing coverage tracked local reductions in sequence complexity, highlighting potential biases associated with difficult-to-map regions.

**Figure 5.**
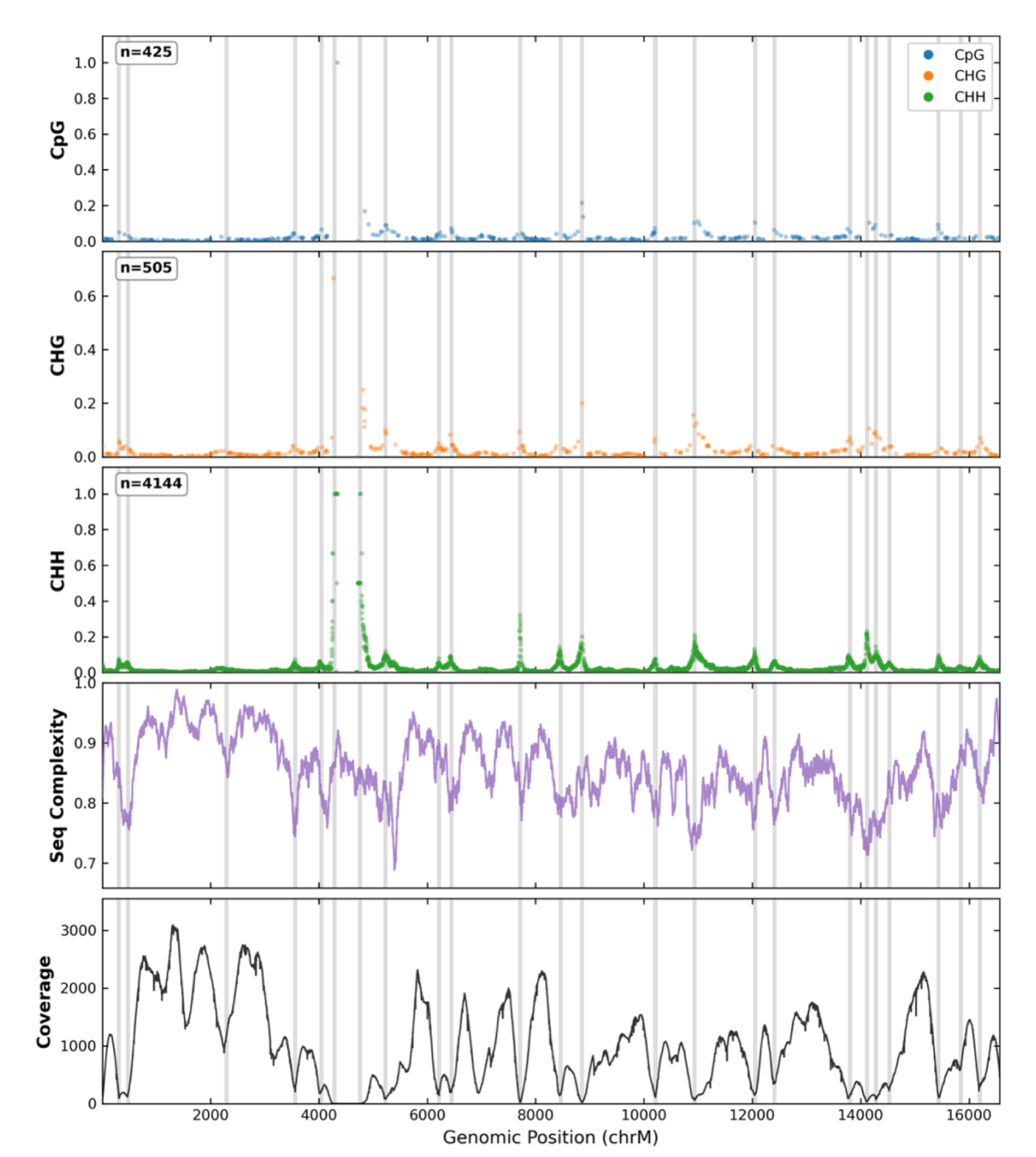
Apparent mtDNA methylation signals on the light strand correlate with low sequencing coverage and reduced sequence complexity. Genomic profiles of 5mC cytosine modification frequencies across the human mitochondrial genome (chrM) light strand. The top three panels display methylation frequencies at CpG (blue), CHG (orange), and CHH (green) contexts. The number of interrogated cytosine sites per context is shown in the top-left corner of each methylation panel. The fourth panel shows normalized sequence complexity (Shannon entropy) computed in 200-bp sliding windows across the reference sequence. Sequence complexity was normalized to a 0-1 scale, where higher values represent greater nucleotide diversity and a more uniform distribution of bases (A, T, G) within each sliding window. The bottom panel indicates per-base sequencing coverage. Vertical gray bars highlight genomic regions with recurrent apparent methylation enrichment across contexts aligned with regions of relative local reduced sequence complexity and lower sequencing depth.

## Discussion

The existence and functional relevance of cytosine methylation in human mitochondrial DNA remain highly debated.^7,10,13,31–33^ Here, we provide convergent evidence from orthogonal sequencing platforms that mtDNA methylation and hydroxymethylation signals are consistent with technical background levels and are influenced by platform-specific and methodological confounding factors. By systematically evaluating mtDNA methylation using enzymatic, native long-read, and short-read sequencing approaches, our findings challenge the interpretation of apparent mtDNA methylation signals as biologically meaningful.

Using these orthogonal approaches, we interrogated the mtDNA across diverse physiological and pathological contexts characterized by elevated or dysregulated nuclear DNA methylation states. These included brain tissue, which exhibits high levels of nuclear DNA methylation and extensive epigenetic regulation^34,35^, stem/progenitor-enriched hematopoietic populations previously reported to display elevated nuclear methylation relative to differentiated cell types^16,35^, and hematological malignancies harboring mutations affecting epigenetic regulatory machinery. Despite these diverse methylation environments, our findings indicate that altered nuclear epigenetic states do not translate into robust or functionally significant mtDNA cytosine modification under the conditions examined.

Locus-specific mtDNA methylation, particularly within the D-loop and mitochondrial coding regions, has been repeatedly reported and interpreted as suggestive of potential regulatory function.^32,36,37^ Several studies have additionally described preferential enrichment of apparent methylation signals on the light strand of the mitochondrial genome^20^. Using Ultima Genomics sequencing and EM-seq, we observed superficially similar strand- and locus-associated patterns; however, these signals were accompanied by pronounced coverage-dependent effects linked to local sequence complexity, suggestive of technical rather than biological signal enrichment. Apparent methylation levels were preferentially elevated on the light strand in regions with reduced sequence complexity, which can challenge short-read alignment and result in reduced coverage relative to average mitochondrial depth. Importantly, some affected loci remained within coverage ranges typically considered high (e.g., >100x), underscoring the limitations of relying on fixed coverage thresholds in mtDNA methylation analyses. Crucially, these observations do not imply that short-read sequencing is inherently unsuitable for mitochondrial methylation analysis. Across the mitochondrial genome, Biomodal sequencing consistently revealed low levels of both 5mC and 5hmC, with no evidence of site-specific enrichment. This approach uses paired-end reads and a chemistry-aware model to assign cytosine modification states, which are encoded separately from the sequence. This enables reconstruction of standard four-base reads with explicit modification annotations and alignment to a four-base reference genome, rather than reduced-complexity (e.g., three-base) genome representations used in many alternative strategies.

By explicitly mapping NUMT-derived CpG sites to their corresponding mitochondrial loci, we demonstrate that nuclear mitochondrial sequences can potentially contribute methylation-like signals across a substantial fraction of the mitochondrial genome. This effect is particularly relevant in antibody-based 5mC/5hmC enrichment workflows applied after mitochondrial isolation, where residual nuclear DNA - including NUMTs - may persist and be enriched during immunocapture. When such enrichment strategies are combined with short-read sequencing and alignment approaches that rely on three-base genomes, ambiguity in read origin can be further amplified, increasing the likelihood of misattribution of nuclear-derived methylation signals to mitochondrial loci. Long-read sequencing provides improved discrimination between mitochondrial and NUMT-derived reads; however, inferred mtDNA modification levels remained sensitive to flow cell chemistry and associated error structures. Notably, in our study an earlier nanopore flow cell chemistry (R9.4.1) yielded higher apparent modification levels than newer chemistry (R10.4.1), which may partially account for previously reported low-level mtDNA modification signals in studies using earlier nanopore flow cells.^19^

We acknowledge that this study is not sufficiently powered for site-level differential methylation analyses due to its sample size. Consequently, we cannot exclude the possibility that rare or stochastic cytosine modifications may occur at isolated sites or within highly restricted mitochondrial subpopulations. However, if present, such events would be expected to occur at levels insufficient to support a broad regulatory role in mitochondrial gene expression or disease biology. We further note that mtDNA cytosine modifications cannot be excluded in certain biological contexts not examined here. In particular, embryonic tissues undergo extensive metabolic and epigenetic reprogramming and therefore remain plausible settings for transient or low-abundance modifications. Beyond physiological conditions, experimentally induced or engineered mtDNA cytosine modifications may also be possible and could provide a useful framework for probing potential mechanisms of mitochondrial gene regulation.

Collectively, these findings indicate that cytosine 5mC and 5hmC modifications are not a detectable feature of mtDNA in adult human tissues under the conditions examined. The combined evidence presented here, together with prior studies reporting a lack of convincing support for biologically meaningful mtDNA methylation,^13,31,33^ suggests that the absence of mtDNA cytosine methylation should represent the default interpretation in human tissues. Accordingly, future studies investigating potential mitochondrial epigenetic modifications will require rigorous methodological and analytical frameworks capable of distinguishing low-abundance biological signal from technical and computational confounding. This includes orthogonal validation of reproducible site-specific signals, systematic assessment of strand bias, local sequence complexity, coverage-dependent effects, and stringent deconvolution of nuclear and NUMT-derived reads.

## Methods

### Patient samples

Acute myeloid leukemia samples were obtained at the time of initial diagnosis, with all patients providing written informed consent for blood collection as part of the University Health Network Hematologic Malignancy Tissue Bank (REB 01-0573). A mobilized peripheral blood stem cell sample was also collected and analyzed as a non-malignant reference sample. All glioblastoma samples correspond to primary tumors, as detailed in Table S2. Informed consent was obtained at St. Michael’s Hospital Brain tissue biobank (Unity Health REB 13-141; University of Toronto REB 39640).

### Biomodal sequencing library preparation and read processing

Genomic DNA libraries representing normal (non-malignant) samples were prepared from brain tissue (BioChain, Catalog No. D1234035) and from blood (lymphocyte cell line NA12878) using the Biomodal kit, following the manufacturer’s protocol. Whole-genome sequencing was performed on an Illumina NovaSeq platform, generating 150 bp paired-end reads. Raw paired-end sequencing reads were trimmed to remove adapter contamination within either read of a pair. The first base of each trimmed read, representing a residual overhang from library preparation, was removed. Reads were subsequently resolved into canonical sequence representations with associated cytosine modification annotations, enabling discrimination of 5-methylcytosine (5mC) in both CpG and non-CpG contexts, as well as 5-hydroxymethylcytosine (5hmC) at CpG sites, using previously described analytical principles.^14^ Reads that were too short, contained excessive ambiguous bases, or exhibited a high proportion of low-quality base calls were excluded from further analysis using custom scripts. Resolved single-strand reads were aligned to the human reference genome (GRCh38/hg38 assembly). The average sequencing coverage across the nuclear genome was approximately 34x for both samples. The read coverage in the mitochondria DNA (mtDNA) was around 51,000× in the brain sample and 20,000× in the blood cell line.

### Differentially methylated region (DMR) calling and data visualization

Per-base methylation information, encoded within custom quality strings, was extracted from aligned BAM files using Samtools.^38^ The mpileup output was then piped to custom AWK scripts specifically designed to parse this encoding. A custom Python script was employed to: i) partition the genome into mtDNA-sized windows; ii) calculate windowed average methylation levels for 5mC and 5hmC; iii) perform statistical comparisons of methylation levels between the brain and blood samples for each window; and iv) compute fold changes. All statistical tests were conducted as two-sided. The window corresponding to the mtDNA was not differentially methylated. Genome-wide visualizations of differential methylation were generated using the karyoploteR package.^39^ Methylation signals, initially encoded in read quality strings, were converted into SAM base modification format using custom Bash scripts. This process generated alignment files containing MM:Z: (modification status) and ML:B:C: (likelihood) tags, suitable for visualizing methylation in IGV.44. In addition, two DMR callers were applied to the mtDNA data: Metilene (v0.2-9)^40^ and Dispersion Shrinkage for Sequencing (DSS; v2.58.0).^41^ Analyses were performed on CpG and non-CpG sites either jointly or separately for 5mC. For 5hmC, only CpG sites were included, as 5hmC calling in this context is restricted to CpG dinucleotides using the Biomodal sequencing framework. Both tools were used for DMR calling with equivalent parameters including requiring a minimum of 10 cytosines per DMR and a minimum 10% mean methylation difference between groups. For Metilene, a maximum gap of 300bp between consecutive positions was allowed within a DMR, while for DSS, DMRs within 300 bp of each other were merged into a single DMR. For DSS, local smoothing was applied to neighbouring cytosine sites to estimate variance.

### Comparison of mtDNA methylation to matched nuclear trinucleotide sequence contexts

To assess mtDNA methylation relative to a comparable nuclear background, we implemented a sampling strategy that controlled for both sequence context and sequencing depth. For each of the 7,162 cytosines located in non-soft-masked regions of mtDNA, analyzed on both the heavy and light strands in brain tissue, we identified the trinucleotide context (5’-NCN-3’), resulting in 16 distinct sequence signatures. For each mtDNA cytosine, we then identified corresponding loci within the nuclear genome of the blood sample that shared the identical trinucleotide signature. Since mtDNA coverage significantly exceeds nuclear coverage (∼51,000X vs ∼34X average), direct comparison of methylation levels is inappropriate. Therefore, to match the high coverage of each mtDNA locus, we randomly sampled multiple nuclear loci sharing the same trinucleotide context until the cumulative read coverage across the sampled nuclear loci approximated the coverage observed at the single mtDNA locus in the brain sample. An aggregate methylation level for this sampled nuclear background was then calculated by summing the methylated base counts across all sampled nuclear positions and dividing by the total number of base counts across these positions.

### Oxford Nanopore sequencing and data analysis

2µg of native genomic DNA extracted from each of three GBM tumor tissues and three AML samples underwent library preparation following the SQK-LSK114 protocol (Oxford Nanopore Technologies). Sequencing was performed using R10.4.1 flow cells. Base calling, including simultaneous 5mC and 5hmC detection, was executed using the Dorado software [https://github.com/nanoporetech/dorado] via the wf-basecalling workflow (version v1.4.5) with the basecalling model and modification model set to dna_r10.4.1_e8.2_400bps_hac@v5.0.0 and 5mCG_5hmCG@v1, respectively. This workflow aligned reads to the GRCh38/hg38 reference genome using minimap2^42^ with soft-clipping enforced (-Y parameter) to enable modification analysis on supplementary alignments. Methylation signals were subsequently summarized per CpG site using Modkit pileup [https://github.com/nanoporetech/modkit]. Analysis was restricted to CpG dinucleotides. The maximum confidence thresholds were required for modification calls. That is, minimum probability > 0.99 for both 5mC (--mod-thresholds m:0.99) and 5hmC (--mod-thresholds h:0.99). Only bases where the probability of the base being correctly called as a cytosine exceeded 0.8 were considered (--filter-threshold C:0.8). Additionally, DNA from GBM patient 1 was also sequenced using the R9.4.1 flow cell and analyzed with an adjusted workflow that preserved the same stringency in modification calling, while employing base calling and methylation models set for the R9.4.1 chemistry.

### Analysis of highly homologous NUMTs to mitochondrial DNA

To identify NUMTs that retain ancestral mitochondrial CpG patterns with very high sequence similarity, NUMT sequences from Wei W, et al,^30^ were aligned against the human mitochondrial reference genome (chrM, hg38 build). For each NUMT, both the forward sequence and its reverse complement were aligned independently using global alignment (pairwiseAlignment function, Biostrings R package) with affine gap penalties (gap opening = 10, gap extension = 1). The alignment orientation (forward or reverse complement) yielding fewer or longer contiguous aligned segments was preferentially selected, provided the longest contiguous segment exceeded 50% of the original NUMT length or was longer than 100 bp. Within the preferred alignment, CpG dinucleotides conserved between chrM and the NUMT were identified. Only NUMTs exhibiting at least 10 conserved CpG dinucleotides were retained. This approach resulted in the final selection of 93 NUMTs, providing high-confidence matches to 796 mtDNA cytosines. Matching NUMTs and mtDNA sequences were extracted from the reference genome using BEDtools^43^.

### Ultima Genomics sequencing and data analysis

Genomic DNA from mobilized stem cells was processed for enzymatic methyl-seq (EM-seq) using the NEBNext EM-seq Kit (New England Biolabs) according to the manufacturer’s protocol and was sequenced on the Ultima Genomics UG-100 platform. Raw sequencing data were processed through the vendor-provided automated analysis pipeline, which performed base calling, quality control, adapter trimming. Alignment was performed on-instrument using the vendor’s Ultima Aligner (UA), mapping reads to a three-letter genome derived from the GRCh38/hg38 reference assembly. Standard alignment metrics and sequencing quality statistics were generated as part of this workflow reporting average aligned read length of 197 bp. Cytosine methylation frequencies were extracted from the aligned CRAM files using MethylDackel (https://github.com/dpryan79/MethylDackel) across CpG, CHG, and CHH sequence contexts. To resolve strand-specific mitochondrial methylation, each cytosine position was annotated according to the reference mitochondrial genome sequence. Consistent with the positional mapping implemented in the analysis pipeline, genomic coordinates in the bedGraph file corresponding to guanine (G) residues in the reference sequence were assigned to the Heavy strand, whereas coordinates corresponding to cytosine (C) residues were assigned to the Light strand.

### Coverage-stratified decay modeling and error thresholding

The relationship between sequencing depth and apparent methylation frequency was modeled using a non-linear exponential decay function: *y* = *a* ⋅ *e*^−*bx*^ + *c*, where x represents coverage, y is methylation frequency, and a, b, and c are fitted parameters. The model was constrained to biologically plausible bounds (0 ≤ *a*, *b*, *c* ≤ 1) and fitted using least-squares optimization. The asymptotic parameter (c) represents the residual background at infinite coverage. Model goodness-of-fit was evaluated using the coefficient of determination (R^2^).

### Sequence complexity

Genomic sequence complexity was computed using a one-base sliding-window approach across the reference mitochondrial genome. A window size of 200 bp was used to approximate the average sequencing read length. To accurately reflect the reads’ sequence complexity following EM-seq, cytosine based were computationally converted to thymine bases prior to calculations, leaving a ternary alphabet (A, T, G). Normalized Shannon entropy was calculated for each window as *H* = −∑*p*_*i*_log _%_(*p*_$_) where *p*_$_ represents the frequency of each base. Entropy values were normalized by the maximum theoretical entropy for a three-base system (log _%_(3) ≈ 1.585), yielding a complexity score ranging from 0 (homopolymeric) to 1 (maximum complexity).

## Supplementary Information

Supplementary Figures - Includes Figures S1-S4

Supplementary Table S1 - List of genomic windows exhibiting differential 5mC and 5hmC enrichment between brain and blood samples.

Supplementary Table S2 - Clinical and genetic characteristics of glioblastoma (GBM) and acute myeloid leukemia (AML) patient samples

Supplementary Table S3 - Summary of mtDNA 5mC and 5hmC data across patient samples and amplified DNA control

Supplementary Table S4 - Detailed information concerning methylation at mitochondrial CpG sites versus their corresponding nuclear-integrated mitochondrial sequences.

Supplementary Table S5: Per-base methylation levels and sequencing coverage in the heavy and light strands of a mobilized blood sample

## Acknowledgements

We thank the patients for their invaluable contributions to this study. We also acknowledge the Ontario Institute for Cancer Research Information Technology Core Facility for providing high-performance computing resources essential to our analyses.

## Authors’ contribution

S.B. conducted computational analyses, developed custom code, curated the accompanying GitHub repository and helped with manuscript writing. P.C.Z. performed sample preparation and Nanopore sequencing for glioblastoma (GBM) tissues under the supervision of J.T.S. D.T contributed to the downstream analyses of data generated using Biomodal and Ultima Genomics sequencing and helped with manuscript writing. A.T.B contributed to the processing and analysis of GBM data under the supervision of J.R. S.D and J.R coordinated the acquisition of GBM samples and related data collection. J.T.S. and A.T.B analyzed the GBM datasets. J.T.S provided expertise in Nanopore sequencing analytics. A.B., T.C., and K.N. performed sample preparation and generated the enzymatic methylation sequencing data under the supervision of B.L. A.A. and M.D.M. coordinated AML patient sample collection and processing. B.L coordinated the acquisition of normal brain and blood samples. S.A. conceived and supervised the study, developed custom code, led the analysis, and wrote the manuscript. All authors contributed to the interpretation of the results and reviewed the manuscript.

## Data availability

All data required to reproduce the results of this study are available via Zenodo at: https://doi.org/10.5281/zenodo.15801168. Custom analysis scripts and figure-generation code are publicly accessible on GitHub at: https://github.com/abelson-lab/mtDNA-Methylation-Revisited. Additional next-generation sequencing datasets not directly supporting the conclusions of this study are available from the corresponding author upon reasonable request and subject to institutional and ethical approvals.

## Supplementary Figures

**Figure S1:**
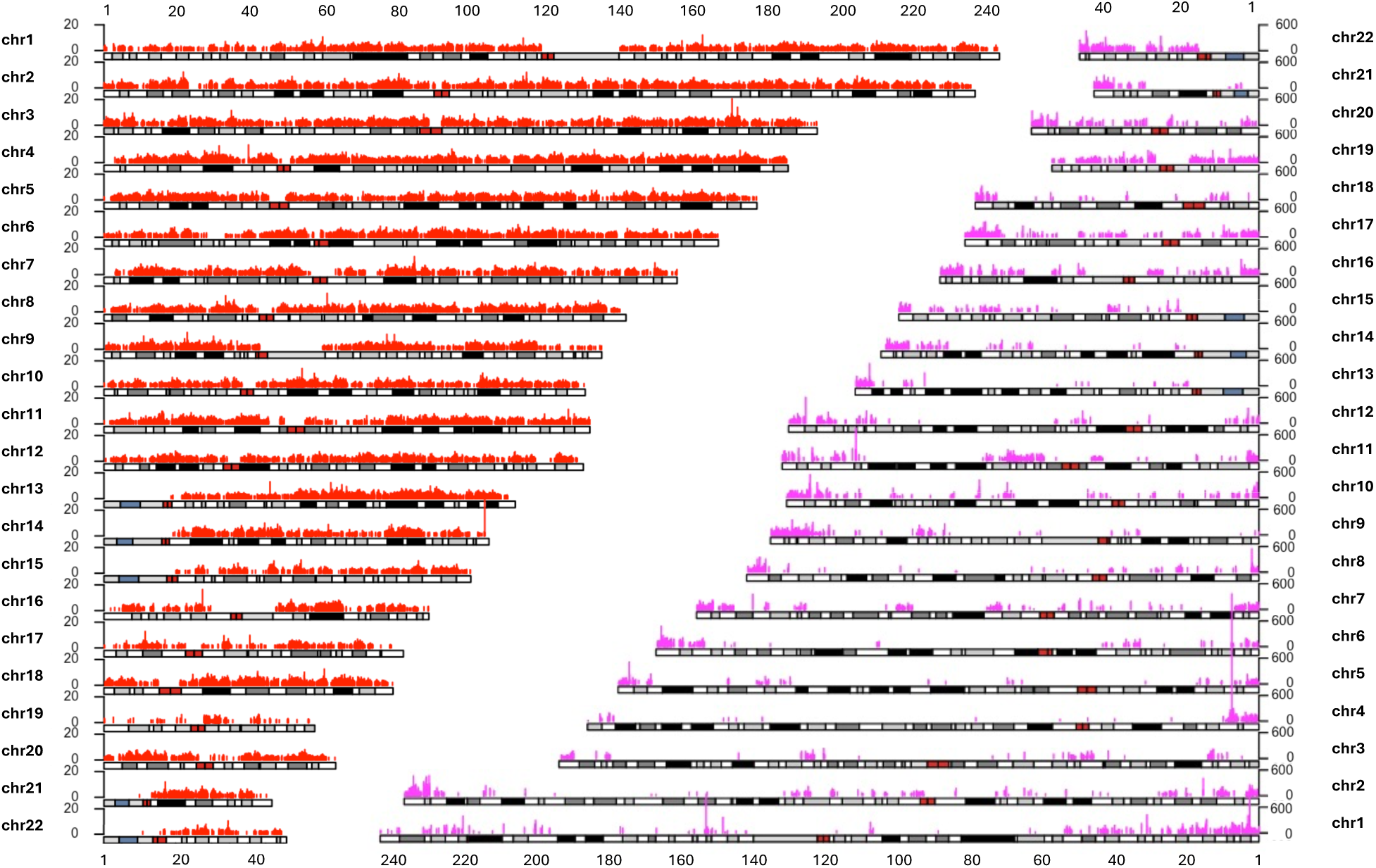
Genome-wide distribution of regions significantly hypermethylated in brain compared to blood. Chromosome ideograms highlight genomic windows with elevated levels of 5mC (left, red) and 5hmC (right, pink). Significance was determined using a Bonferroni-corrected p-value threshold < 0.05 and a Brain/Blood fold change >= 2. Bar heights represent the fold change, indicating the magnitude by which brain methylation exceeds blood methylation in these specific loci. Y-axes are scaled to maximum fold changes of 20 for 5mC and 600 for 5hmC. Black and dark grey bands represent heterochromatin regions while light grey and white bands indicate euchromatin regions.

**Figure S2:**
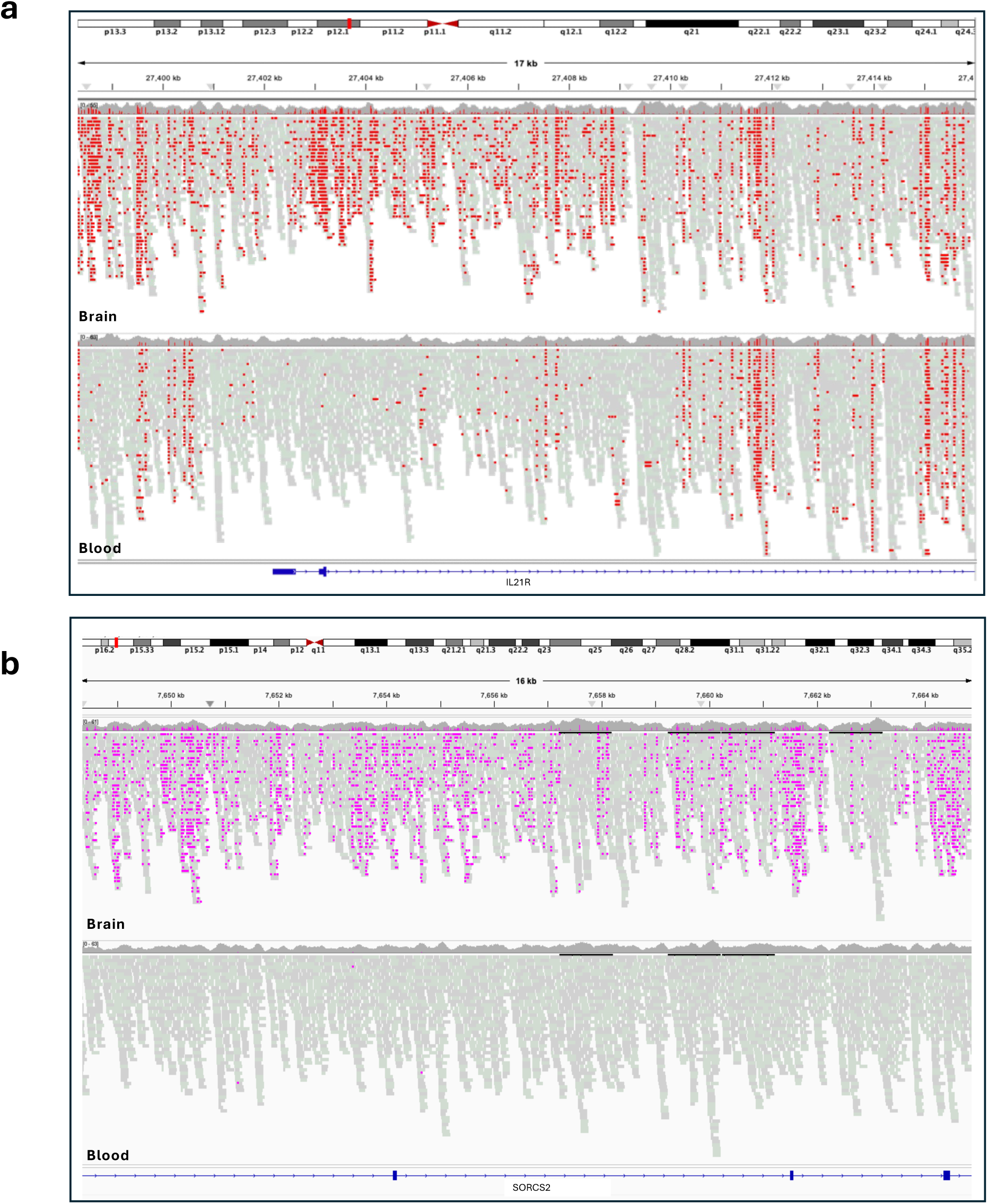
Integrated Genomics Viewer (IGV) snapshots representing regions with high fold-changes for 5mC and 5hmC in brain compared to blood, highlighting tissue-specific epigenetic patterns consistent with known gene functions. **a**. Decreased 5mC at CpGs of the Interleukin-21 receptor gene, showing 13-fold lower levels in the blood sample relative to brain. **b**. Increased 5hmC at CpG sites within the SORCS2 gene, exhibiting 2,974-fold higher levels in the brain sample.

**Figure S3:**
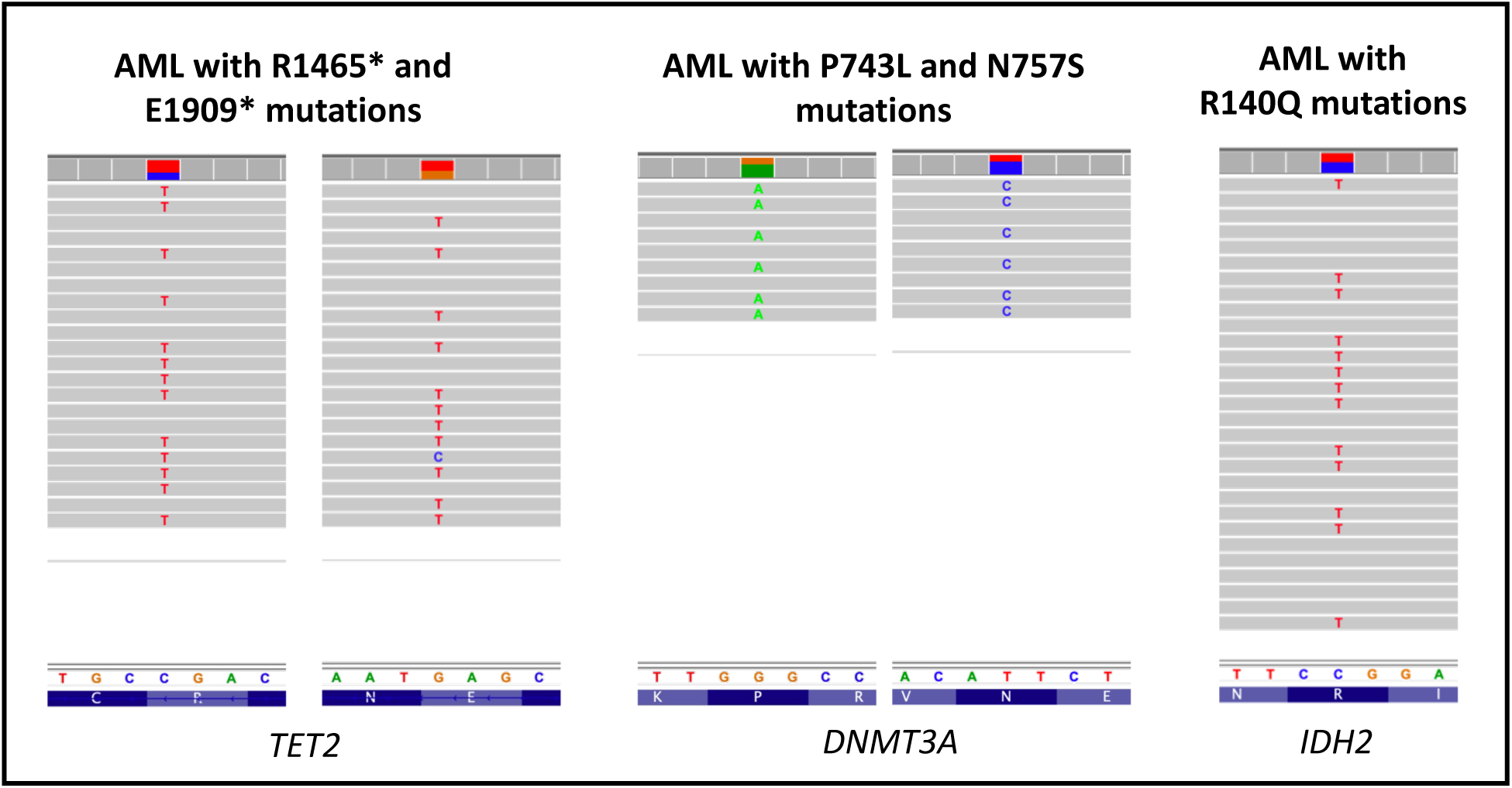
Nanopore sequencing validation of mutations identified during clinical sequencing of acute myeloid leukemia patients. Integrated Genomics Viewer (IGV) snapshots display base substitutions corresponding to five distinct somatic mutations detected across three patients.

**Figure S4:**
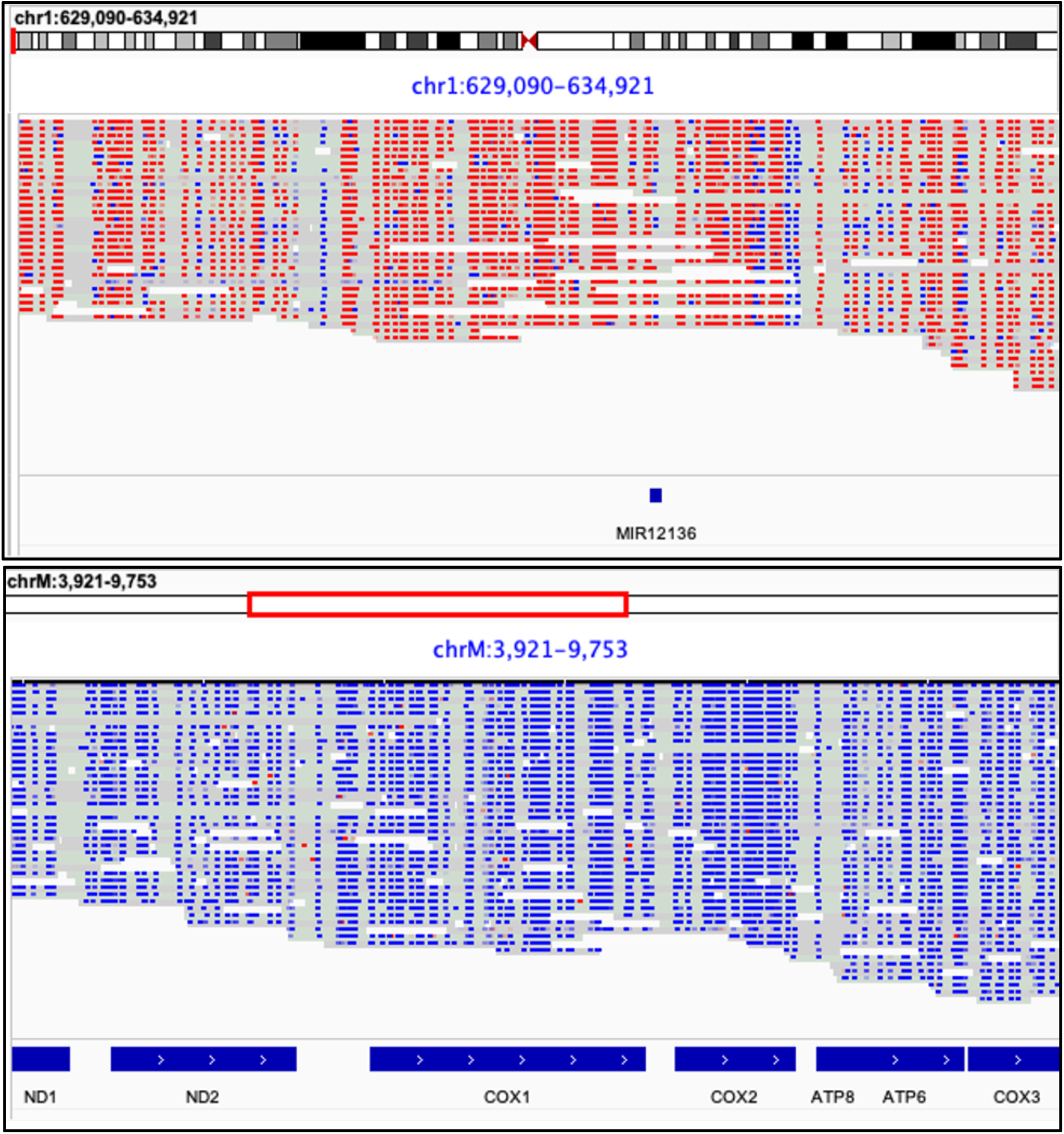
Contrasting methylation landscapes of NUMTs and homologous chrM. IGV snapshot illustrate methylation patterns derived from nanopore sequencing data for sample GBM patient 1. Upper panel shows specific NUMT loci (genomic coordinates indicated). Lower panel shows the corresponding ancestral mtDNA region. Individual CpG methylation predictions on nanopore reads are visualized: red for methylated, blue for unmethylated. The example highlights consistent 5mC within these nuclear-integrated sequences versus the general absence of predicted methylation signal in their mitochondrial counterparts.

